# GAMBIT (Genomic Approximation Method for Bacterial Identification and Tracking): A methodology to rapidly leverage whole genome sequencing of bacterial isolates for clinical identification

**DOI:** 10.1101/2022.06.14.496173

**Authors:** Jared Lumpe, Lynette Gumbleton, Andrew Gorzalski, Kevin Libuit, Vici Varghese, Tyler Lloyd, Farid Tadros, Tyler Arsimendi, Eileen Wagner, Craig Stephens, Joel Sevinsky, David Hess, Mark Pandori

## Abstract

Whole genome sequencing of clinical bacterial isolates has the potential to transform the fields of medicine and public health, with particular impact on molecular epidemiology, infection control and assessing the spread of antibiotic resistance. To realize this potential, bioinformatic software needs to be developed that meets the quality standards of a diagnostic test to allow the reporting of identification results. Our research group has developed a methodology (GAMBIT: Genomic Approximation Method for Bacterial Identification and Tracking) using k-mer based strategies for identification of bacteria based on whole genome sequence reads. GAMBIT incorporates this algorithm with a highly curated searchable database of genomes, which is a subset of the NCBI RefSeq assembly database. In this manuscript, we describe the validation of the scoring methodology, robustness to chosen parameters, establishment of confidence thresholds and the curation of the reference database. We discuss a validation set with GAMBIT deployed as a laboratory-developed test at the Alameda County Public Health Laboratory in Oakland, California. Three advancements were required to build upon existing k-mer based strategies to allow GAMBIT to possess the quality control parameters desired for its use as a diagnostic laboratory-developed test. Firstly, we innovated the data structure used to store the database of known bacterial genomes. This allowed us to store 48,224 bacterial genomes in a k-mer based database—a majority of the bacterial genomes in the NCBI RefSeq database at the time of development. Secondly, curation of the NCBI RefSeq database was required to remove ambiguous or potentially incorrectly labeled bacterial genomes to greatly increase confidence in positive matches. Lastly, we used this curated version of the NCBI RefSeq database and our scoring method to generate confidence thresholds for identification. Thus, the end-user does not rely simply on the closest match, but is informed whether that closest match exceeds a threshold for highly confident identification. This method greatly reduces or eliminates false identifications which are often detrimental in a clinical setting.

## Introduction

Bacterial species that cause human disease are an ever evolving threat and a leading cause of death world-wide^1^. Because of this evolving threat, identification and characterization of these pathogens are crucial for effective infection control methods^2-3^. The potential of whole-genome sequencing (WGS) of clinical isolates to rapidly identify bacterial species, detect antibiotic resistance and provide relatedness information for epidemiological purposes is broadly recognized in the field of clinical microbiology^4-7^. Molecular diagnostic tools based on nucleic acid identification have long been used in clinical microbiology. Clinical microbiology represents approximately 70% of the global market for molecular diagnostic tools, so the extension of WGS as a diagnostic tool is a natural fit^8-9^. WGS has been deployed in clinical microbiology labs as proof-of-concept, used for surveillance of antibiotic resistance or focused on a specific genus or species of clinical relevance, but the literature does not detail the routine use of WGS by clinical microbiology laboratories for identification of bacterial isolates^7,10-11^.

Over the past decade, improvements to Next Generation Sequencing (NGS) have dramatically increased access to draft genome sequences (especially in bacteria) and have substantially decreased in cost per genome^10,12^. This technology has led to expanded use of whole genome sequencing (WGS) in microbial research^13-14^. This research has produced expanded insights into microbial diversity, antibiotic resistance^15-16^, the human microbiome and the evolution and spread of microbial pathogens^17-19^.

Several problems have been noted in the literature that prevent routine use of WGS for clinical microbiology. Consistent among reviews on the topic is that bioinformatic pipelines are needed that process the genomic data once generated^10-11,20^. As Besser^10^ notes, “It is a challenge that the comparability of the sequence data generated on different platforms with different error profiles using different library preparation methods has still not been comprehensively assessed and validated.” Our methodology is agnostic to the platform on which the raw sequence data was generated and allowed our reference database to include over 48,000 bacterial genomes generated from a variety of whole genome sequencing methods. In this manuscript we describe a methodology to rapidly compare a single new bacterial genome sequence to a reference database of over 48,000 bacterial isolates containing 1,415 species. We describe our approach to defining conservative thresholds that make false identification of a bacterial isolate highly unlikely. We demonstrate this algorithm performs well over a variety of test sets. Furthermore, we detail the deployment of this methodology in a clinical setting and describe the results against 88 meticulously identified samples received by the Laboratory over a three year period. The results presented here demonstrate that our methodology provides the necessary bioinformatic elements necessary for use of WGS in a clinical microbiology laboratory.

## Materials and Methods

### Obtaining clinical samples from Alameda County Public Health Labs

Bacterial organisms were obtained and selected in either of two ways: a) isolates generated at the Alameda County Public Health Laboratory through normal diagnostic requests from jurisdictional hospitals were identified through validated and FDA cleared methods. The primary method was API Strip (Biomerieux, Durham, NC). Isolates so identified were confirmed for genus and species identity by reference laboratory (California Department of Public Health) using the same or alternative methods; another means, b) included the use of bacterial isolates provided by proficiency testing (College of American Pathologists). Isolates received for quality assessment were saved by freezing the acquired isolates and subsequently analyzed through whole genome sequencing and then by GAMBIT. Consensus identification was verified by report from the College of American Pathologist and compared to GAMBIT results.

### Obtaining clinical samples from Nevada State Public Health Labs

Bacterial organisms were obtained from routine specimen submissions at the Nevada State Public Health Lab. The organisms were identified through validated and FDA cleared methods. Most bacterial identification was performed using BD Phoenix M50 and MALDI-TOF (Siruis, Bruker). Mycobacterium were identified using AccuProbe (Hologic). Additional species identification was performed on enteric bacteria using ANI (Average Nucleotide Identity) in BioNumerics 7.6 (Applied Maths).

### Whole genome sequencing from Alameda County Public Health Labs

Isolates were collected from culture plates and were subject to genomic DNA extraction using the MagNAPure Compact (BioRad, Hercules CA). The Bacterial lysis procedure was used, according to the package insert. Extracted genomes were assessed for DNA concentration by Qbit fluorescent analyzer. Library preparation was performed by using the Nextera XT (Illumina, La Jolla, CA) with the following alteration to the package insert protocol: final library quantitative normalization was not performed by using the bead-based method in the procedure. Instead, final library concentrations per specimen were determined by fluorimetry and were equalized by dilution in water. Sequencing included single-end read, using 2×150 sequencing kits. Final FASTQ files were generated on the MiSeq instrument and were transferred by external hard drive to a computer equipped with bioinformatic software and GAMBIT for final analysis.

### Whole genome sequencing from Nevada State Public Health Labs

Isolates were collected from culture plates and were subject to genomic DNA extraction using DNeasy Blood and Tissue kit on a QiaCube instrument (Qiagen). Extracted genomes were assessed for DNA concentration by Qubit 3 Fluorometer (Invitrogen). Library preparation was performed by using the Illumina DNA Prep kit (Illumina). Paired-end sequencing was performed, using 2×150 MiniSeq Mid-Output sequencing kits (Illumina). Final FASTQ files were generated on the MiniSeq instrument and were transferred through the cloud to Terra.bio for GAMBIT analysis.

### GAMBIT signature generation and genomic distance metric

A GAMBIT signature is a compressed representation of a genome sequence which supports efficient calculation of the GAMBIT genomic distance metric. It is defined as the set of k-mers present in the genome which occur immediately following a fixed prefix sequence. The signature depends on the value of two parameters - k (a positive integer) and the prefix. The genome database GAMBIT uses for taxonomic classification uses the values *k* = 11 and prefix = ATGAC. The software supports calculation of signatures from genome assembly files in FASTA format.

Internally, the GAMBIT software library represents k-mers using integer indices in the range[0 . . 4^*k*^ − 1]. Indices are derived using the alphabetical ordering of all 4^*k*^ k-mers for the given value of *k*. Thus the 5-kmer AAAAA has index 0 and TTTTT has index 4^5^ − 1 = 1023. Full genome signatures are stored as arrays of k-mer indices in sorted order using the smallest possible unsigned integer data type. Reference genome signatures in the GAMBIT database consist of 7,000 32-bit integers on average, requiring only 28 kilobytes of storage. The software defines a binary file format which is capable of storing any number of genome signatures alongside arbitrary metadata. The GAMBIT distance between two genomes is calculated as the Jaccard distance between the k-mer sets that comprise their signatures. Naturally, the two signatures must have been calculated using identical parameter values (*k* and prefix sequence).

The GAMBIT signature is analogous to the concept of a “sketch” in the Mash tool^21,23^ in that it is a representation of the k-mer content of a genome which supports calculation of a genomic distance metric based on the Jaccard distance. The key difference is in how the two tools accomplish this task with a level of computational efficiency that supports running tens or hundreds of thousands of such comparisons in a short time on modest hardware. Mash uses the MinHash^24^ technique to create a highly compressed “sketch” of the full k-mer content of a genome, which can be used to approximate the true Jaccard distance between the sets represented by two such sketches. By contrast GAMBIT performs a subsampling of the genome’s full k-mer set and calculates the Jaccard distance between two of these subsets directly. Despite its relative simplicity, we show genomic distances calculated using our method still correlate very highly to ANI.

### GAMBIT taxonomic classification

The GAMBIT database used for classification consists of precalculated signatures for 48,224 reference genomes along with additional genome metadata and a set of taxonomy nodes. Taxonomy information is derived from the NCBI taxonomy database, but restricted to the genus and species ranks and subject to additional manual curation. Each reference genome is assigned to a taxonomy node, typically of species rank. In a small number cases where manual editing of the taxonomy structure was required, genomes were assigned to artificial nodes below the species level. Genus and species nodes are assigned a threshold distance.

The classification process takes as its input an assembled query genome in FASTA file format. GAMBIT first calculates the signature of the query sequence and then the distance from the query to each reference genome signature. From this it selects the reference genome with the minimum distance to the query, and compares this distance to the threshold of the reference genome’s assigned species. If the distance is less than this threshold then GAMBIT will classify the query as the given species. Otherwise GAMBIT will ascend the taxonomy tree from the species node to its parent genus node and repeat the same process, possibly returning a genus-level classification. If the query does not fall within the genome’s threshold distance GAMBIT will be conservative and report the query as “unknown.” Regardless of the classification made, GAMBIT will always report the closest reference genome along with its taxonomic assignment and distance to the query.

Although the current database only makes classifications at the species and genus levels, this scheme supports deeper and more complex taxonomy trees that could be used to make classifications at the subspecies level.

### GAMBIT tree generation

GAMBIT includes a function to estimate a relatedness tree for a given set of genome assemblies. It calculates pairwise distances between all input genomes and then performs hierarchical clustering using the UPGMA method. The resulting tree can be output in Newick format or visualized as a dendrogram.

### ANI score generation

ANI scores in this manuscript were generated using the FastANI tool^22^ with default parameter values.

### Performance benchmarks

Performance of the full GAMBIT classification process was measured on a Dell XPS laptop with a six-core Intel i7-9750H processor and 16 gigabytes of memory, with three replicates run for each genome set 1-4.

## Results

### A k-mer-based representation supports a reference library of over 48,000 bacterial genomes

Whole-genome sequencing (WGS) data have been used for species identification using k-mer based methods^21,23^. A previously described k-mer based strategy of identification^21^ was limited to ∼1,700 reference genomes based on their data storage structures. GAMBIT was able to extend its reference database to 48,224 bacterial genomes by utilizing a different method of representing genomic data. GAMBIT finds all 11-mer sequences in WGS data that immediately follow the prefix sequence ATGAC. There are only 4^11^ = 4,194,304 such sequences. By associating each of these 16-mers with an integer in the range 1 . . 4^11^corresponding to its position in alphabetical order, the set of unique 16-mers present in a genome can be represented with a simple integer array. This storage method allows the entire database of 48,000+ reference genomes to fit in less than 1.4GB, and supports rapid screening of unknown genomes against the full set of references. The entire GAMBIT classification process took an average of less than .5 seconds when run on a personal laptop for all genomes in sets 1-4, suggesting that the process can be scaled to far larger reference databases.

### GAMBIT genomic distance metric correlates with sequence identity

We utilized four sets of bacterial genome assemblies in validating the GAMBIT distance metric (Table 1). The first two consist of high-quality assemblies obtained from the NCBI RefSeq database, while the second two are derived from clinical samples. Set 1 (Supplemental Table 1) consists of 492 *E. coli* genomes used in Ondov *et al* ^23^ to validate the Mash tool, which defines a similar k-mer based genomic distance metric. Set 2 (Supplemental Table 2) is composed of 70 completely closed genomes that broadly cover the bacterial kingdom based on Konstantinidis and Tiedje^25^. Bacterial groups include 15 genomes from *Enterics*, 9 from *Streptococcus*, 7 from *Staphylococcus*, 4 from *Bacillus*, 4 from *Mycobacterium*, 4 from *Neisseria*, 3 from *Bordetella*, 3 from *Pseudomonas* and 21 other genomes representing the following genuses: *Brucella, Burkholderia, Clostridium, Heliobacter, Legionella, Rickettsia, Tropheryma, Vibrio, Xanthomonas* and *Xylella*. Set 3 consists of 88 “gold standard” proficiency test samples, described in detail later in the manuscript. The source of these proficiency test samples is described in the Materials and Methods. They are part of the routine testing for clinical diagnostic labs spanning 46 different species over 28 unique genuses. Lastly, Set 4 is a set of 605 genomes obtained between 2020-2022 by Nevada State Public Health Labs. It contains 25 species and 15 genera.

**Table 1:** List of genome sets used in this manuscript.

| Data Set | Number of Genomes | Source of fasta/fastq files | Phylogenetic Diversity | Assembly Quality | Reference |
| --- | --- | --- | --- | --- | --- |
| Set 1 | 492 | NCBI Ref Seq | Low (*E. coli* only) | Medium | Ondov *et al* (23) |
| Set 2 | 70 | NCBI Ref Seq | High (multiple phyla) | High (all reference genomes) | Konstantinidis and Tiedje (24) |
| Set 3 | 88 | Alameda County Public Health Labs | High (multiple phyla) | Medium | none |
| Set 4 | 605 | Nevada State Public Health Labs | High (multiple phyla) | Medium | none |
| Set 5 | 29 | Stephens Lab | Low (*E. coli* only) | High (all closed genomes) | Stephens *et al* (27) |

We used ANI (Average Nucleotide Identity)^26^ as a baseline measure of genomic similarity to validate the GAMBIT distance metric. ANI is generally used to determine similarity at the species or genus level with thresholds above 0.92 being optimal for species level calls^26^. We compared pairwise ANI values against GAMBIT distances in each of our comparison datasets (Figure 1). Spearman correlation was high for all three sets (Set 1 = 0.977; Set 2 = 0.968; Set 3 = 0.969; Set 4 = 0.978). Of note for Set 4, GAMBIT demonstrates broader discrimination for genome pairs with ANI near 100%, resulting in a number of values off the major trendline.

**Figure 1.**
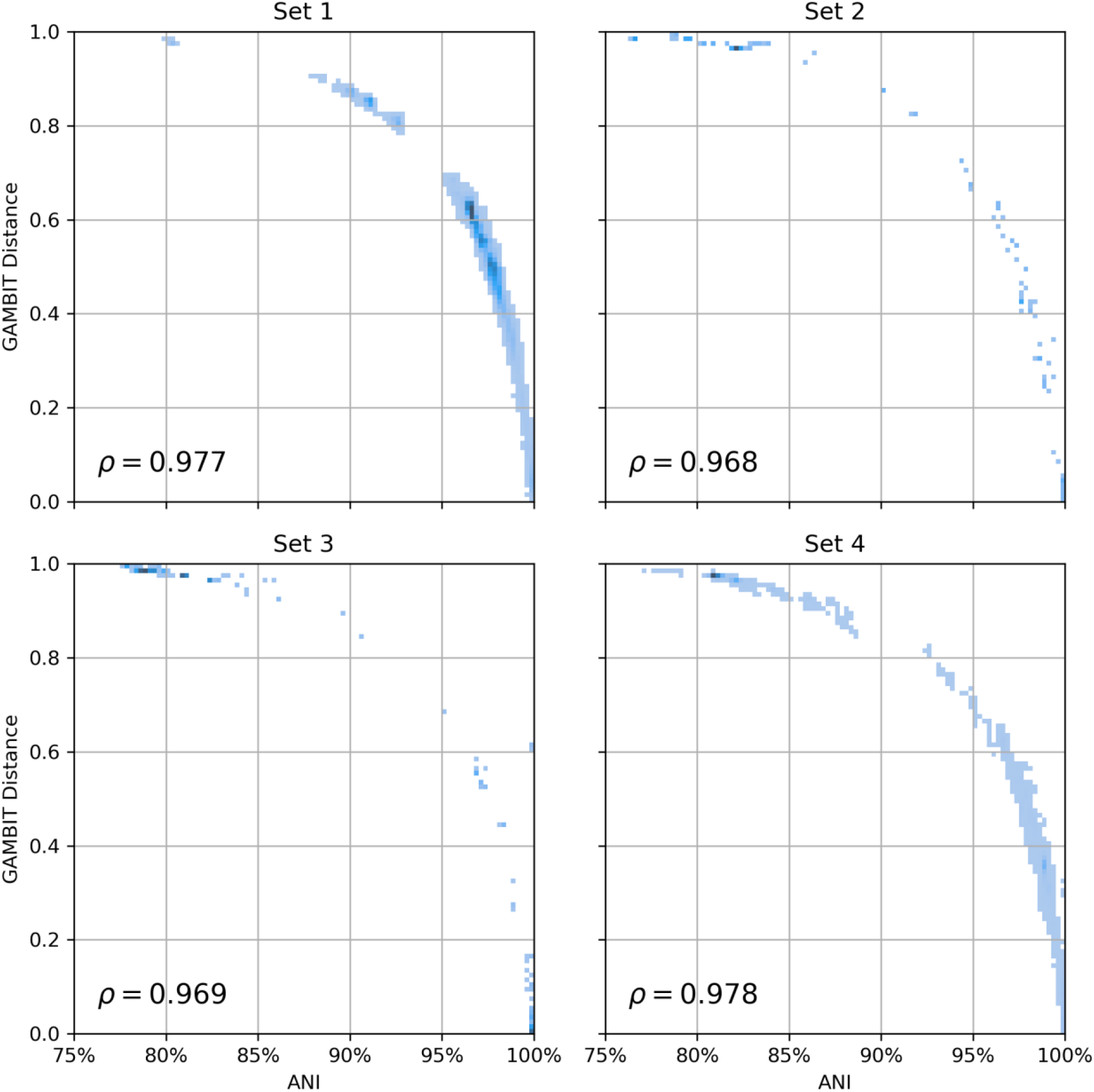
Relationship between GAMBIT distance and ANI (Average Nucleotide Identity) for all pairs of genomes in each data set 1-4. The relationship is nonlinear but close to monotonic as measured by Spearman correlation (absolute value in the bottom left corner of each subplot). ANI was calculated using the FastANI tool (see materials and methods) with default parameter values. GAMBIT distances were calculated using the same parameter values as the RefSeq reference database (k=11, prefix=ATGAC). As FastANI only reports ANI values great than ∼80%, the fraction of total pairwise comparisons shown here were 100%, 5.5%, 7.1% and 47.4% for data sets 1-4 respectively.

### GAMBIT distances are robust with regard to parameter choice

We wanted to test three parameters in the GAMBIT algorithm: the sequence of the anchor prefix used, the length of the k-mer and the length of the anchor sequence. First, we tested the sequence of the anchor. We used ATGAC as our prefix as we based the logic of our algorithm on work from the Aarestrup group^21^ that used the same prefix sequence. We generated seven additional prefix sequences at random of length seven. We tested each of these prefixes varying the lengths of the prefixes from length 4 to length 7. We varied the k-mer length from 7 to 17. In Figure 2 we show results against our test data sets described above and in Table 1. In Figure 2 we show the Spearman correlation between ANI and GAMBIT distance with each data point representing a unique set of GAMBIT parameters. In Figure 2A, the four plots represent different prefix lengths (prefix length 4 on the far left to prefix length 7 on the far right). All panels use the same axes and are comparable. The color of each line represents a different data set. Along the x-axis, k-mer length is varied from 7 to 17.

**Figure 2.**
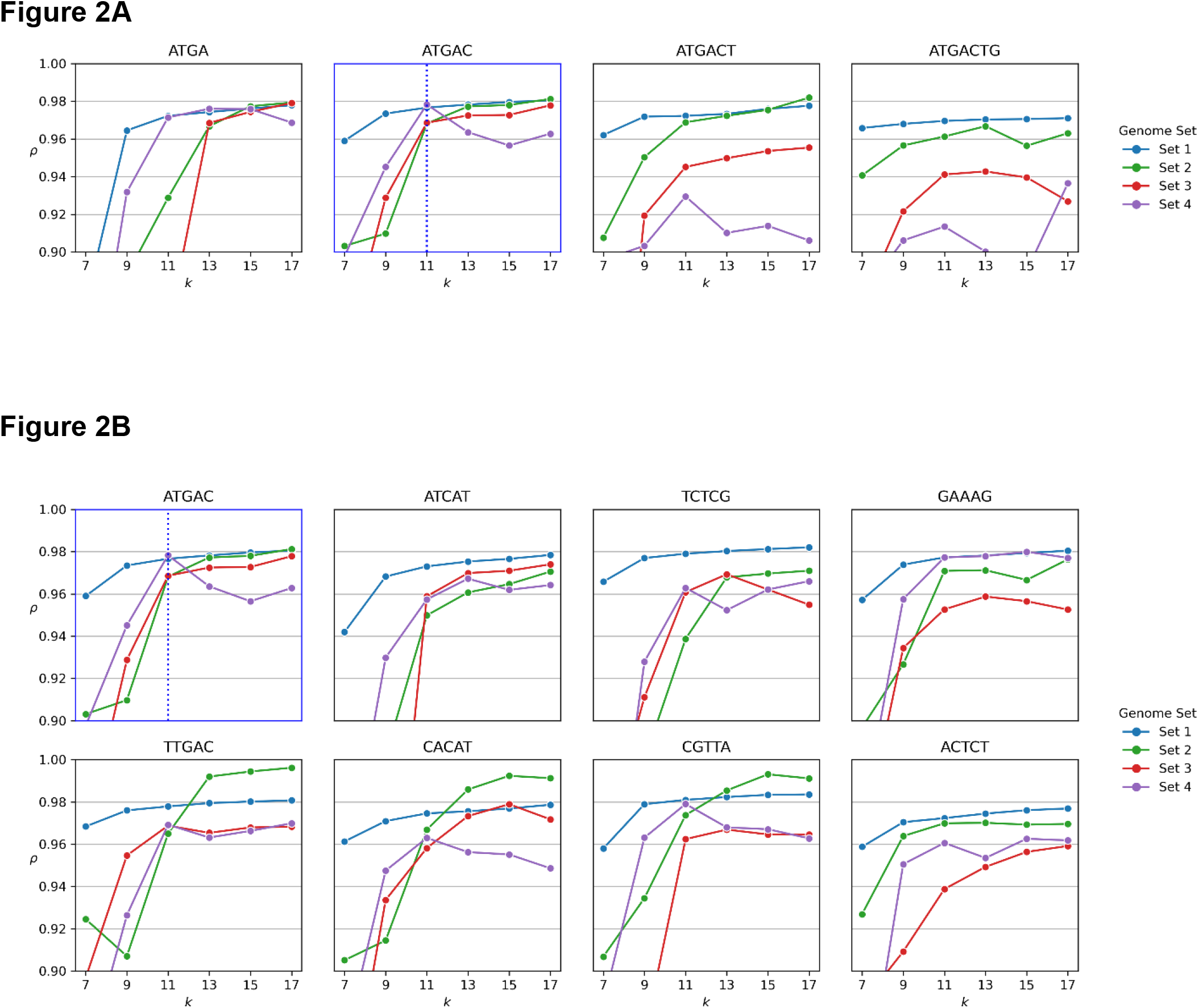
Spearman correlation between GAMBIT distance and ANI for different GAMBIT parameter values. Each subplot in both A and B represents a different choice of prefix sequence. Subplots show the absolute value of the Spearman correlation vs. value of the k parameter for all pairwise comparisons within genome sets 1-4 (Table 1). Standard values of the prefix and k parameters used throughout the rest of the manuscript are highlighted by a blue subplot border and blue vertical line, respectively. A. Variations of our standard prefix ATGAC with length between 4 and 7 nucleotides. B. Standard sequence plus 7 random sequences of the same length.

The set of parameters used in the final version of GAMBIT are shown in Figure 2A on the plot with the blue highlighted border. Beginning with this plot, we would expect to see the highest Spearman correlation in the upper right corner of the plot--which corresponds to the longest length of those k-mers (the longer the k-mers, the more of the genomic information is being retrieved). These data points represent the maximum amount of genomic information being used by the GAMBIT algorithm and they do represent the highest Spearman correlations for each set of variables. In terms of k-mer length, performance increases in all instances as k-mer length increases reaching a plateau at a k-mer length of 11. This suggests that our chosen k-mer length of 11 balances discriminatory power with speed of the algorithm.

Comparing all four panels in Figure 2A, we observe the effect of changing the length of the prefix. We predict that as prefix length increases correlation should decrease because less of the genome is being sampled. Interestingly, correlations do not decrease significantly when increasing the prefix length to 5 (the chosen parameter for our version of GAMBIT), but degradation in performance is seen when the prefix length is increased to 6 or 7 nucleotides.

Lastly, we observe the effect of the actual sequence of the prefix sequence. We compare our prefix sequence (at length 5) to seven other randomly generated prefix sequences (Figure 2B). We compared Spearman correlation between ANI and GAMBIT distance for each sequence over k-mer lengths ranging from 7 to 17. Our chosen prefix sequence (ATGAC), performs in the middle of the different prefix sequences across all comparison sets. Without understanding why specific prefix sequences perform better or worse, using a sequence that is consistently in the middle should minimize the chance of introducing bias into our method based on sequences overrepresented in our comparison dataset but not present broadly among all bacterial species. A more exhaustive comparison of parameters is shown in Supplemental Figure 1.

### Curated, High-Quality Database construction based on RefSeq

There is no large-scale, gold-standard reference database of bacterial genomes. NCBI RefSeq database is the best contender. Our goal in creating a database for GAMBIT was two-fold. First, the genomes representing each species in our database must contain enough distinction from other species that similarity thresholds could be extracted (see next section). Second, that we have the highest level of confidence that a genome in our database was actually from the species that matched its labeled identification. Interrogation of the RefSeq database clearly identified incidents of mislabeled genomes. The conservative set of filters used to achieve our two stated goals are described below.

To construct our reference database, we started with the 60,857 available bacterial genomes from the NCBI RefSeq database^27^ on July 1^st^, 2016. We purged the 3,996 genomes that did not have an associated genus and/or species. Additionally, we purged the 3,305 genomes that were the only representative of its species—so each species must have at least two separate sequenced isolates to appear in our reference database. After the removals noted above, we calculated all pairwise Jaccard scores for the remaining 53,556 genomes. We then began an iterative process of removing ambiguous genomes, resulting in the removal of an additional 5,332 genomes. We defined ambiguous genomes as genomes that met any of the three following criteria: (1) any genome that did not cluster well with the majority of the other genomes within their species, (2) any genome that clustered well with some members of their species but also several members of another species in the database, or (3) any genome that did not cluster well with any genomes in the database. We chose to eliminate these genomes to attain a higher level of certainty in the final database, which contains 48,224 bacterial genomes representing 1,414 species and covering 454 unique genera. The remaining genomes in our reference database represent a large set of genomes over a large breadth of clinically relevant species (genome list in Supplementary Table 3). This gold-standard of bacterial genomes and their genus and species identification should be a valuable resource for other microbial genomic projects.

### The necessity of draft genome assembly for generated fastq files

Analysis of the copy numbers of unique k-mers found in raw sequencing data (Figure 3) reveals a bimodal distribution. This is shown for five different whole-genome sequences selected from Set 3. Histograms are split based on whether k-mers werewere filtered out after assembly (blue) or still included after assembly (green). The high copy number peak consists almost entirely of k-mers retained in the assembly and is centered around the sequence coverage depth (represented by the dashed black vertical lines). This is expected as sequence coverage depth represents the average number of sequence reads that cover a particular sequence, thus the number of times each unique k-mer is identified in a given sequencing experiment should be approximately equal to the sequence coverage depth.

**Figure 3.**
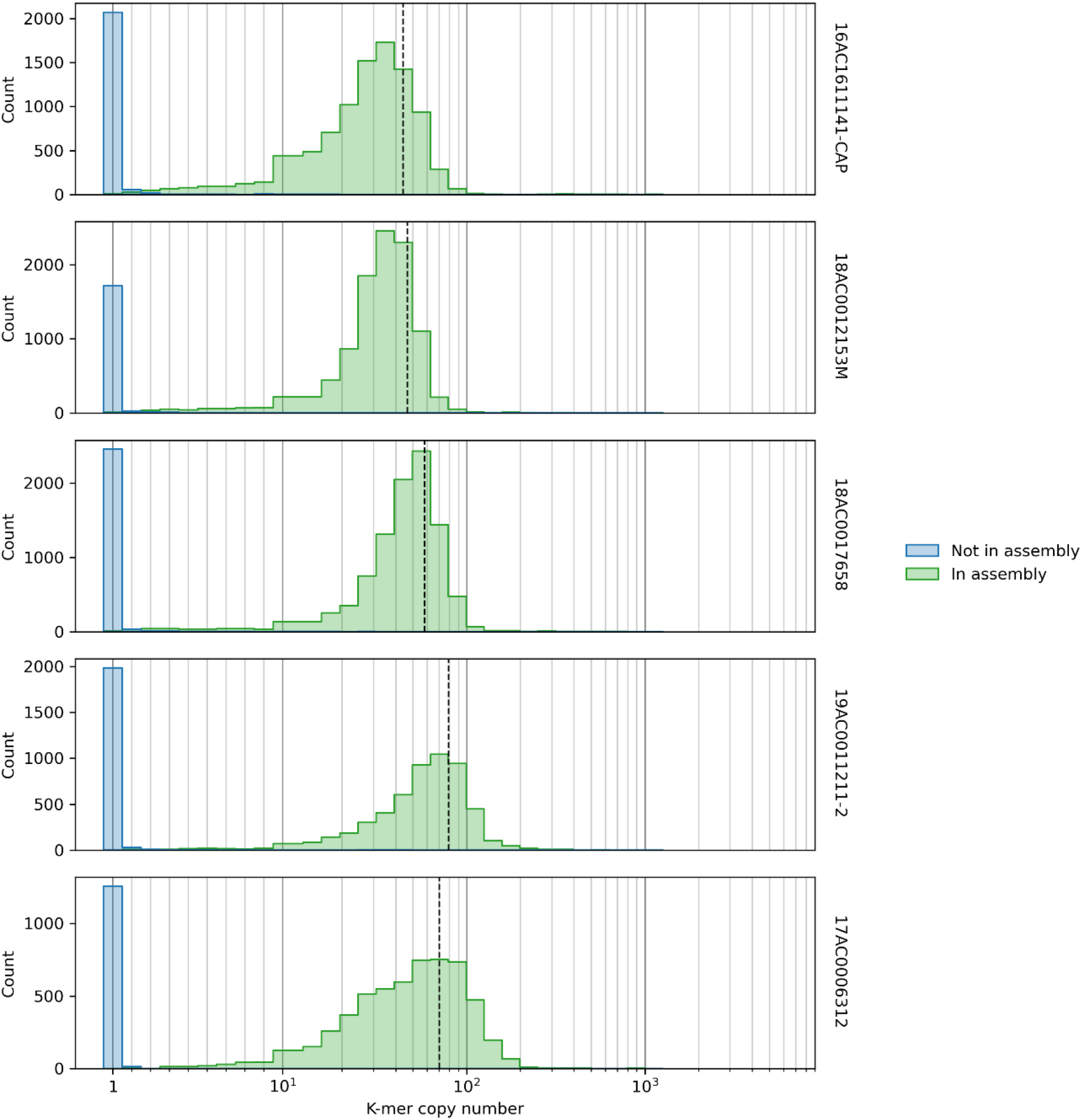
Distribution of k-mer counts in raw sequencing reads. Raw FASTQ files for a selection of Set 3 genomes were scanned for k-mer matches using the standard GAMBIT procedure and the number of copies of each unique k-mer found was tallied. Species: A is *P. stuartii*, B is *E. coli*, C is *Y. enterocoliticia*, D is *E. faecalis* and E is *S. epidermis*. The blue histograms show the copy number of k-mers found in the raw sequencing reads that are not recovered after assembly and likely represent sequencing errors. The green histograms show the copy number of k-mers found in the sequencing reads that are present in the final genome assembly. The center of these distributions closely tracks the sequencing depth (denoted by the black dashed vertical line) found in the assembly of that particular fastq. This is to be expected as the green histogram represents the actual k-mers found in the genome of the organism of interest.

The low copy number peak consists almost entirely of k-mers excluded from the final assembly which were predominantly found only once in the fastq file, with a short right-sided tail. This distribution represents sequencing errors which would account for the vast majority of k-mers in this distribution being found only once and that these k-mers are eliminated after assembly of the fastq file into a fasta file. This is because in generating the consensus sequence in the process of a *de novo* genome assembly nearly all sequencing errors are eliminated. For GAMBIT to work properly, users must perform an assembly of the fastq file to eliminate sequencing errors by consensus.

### Establishing confidence thresholds for generating “resulting identifications” vs. “suggested identifications”

To utilize GAMBIT as a diagnostic test, it was crucial to establish thresholds for definitively calling a particular genome at the species or genus level. For most species, we could use the most conservative threshold which involved calculating two parameters: max intra distance (or diameter) and min inter distance (Figure 4). First, we generate all pairwise comparisons for all isolates of a species within that species and identify the largest distance (we refer to this value as max intra or diameter). Max intra represents the GAMBIT distance between the two most dissimilar isolates within a species in our reference database. Next, we generate all pairwise comparisons between isolates within our species and every other isolate from other species in our database. From those values we would identify the smallest distance (we refer to this value as min inter). Min inter represents the closest similarity that can be generated by another species in our database to any isolate of our species of interest in our database. Put another way, min inter represents the best false positive value that could be generated by the 48,224 genomes in our reference database.

**Figure 4.**
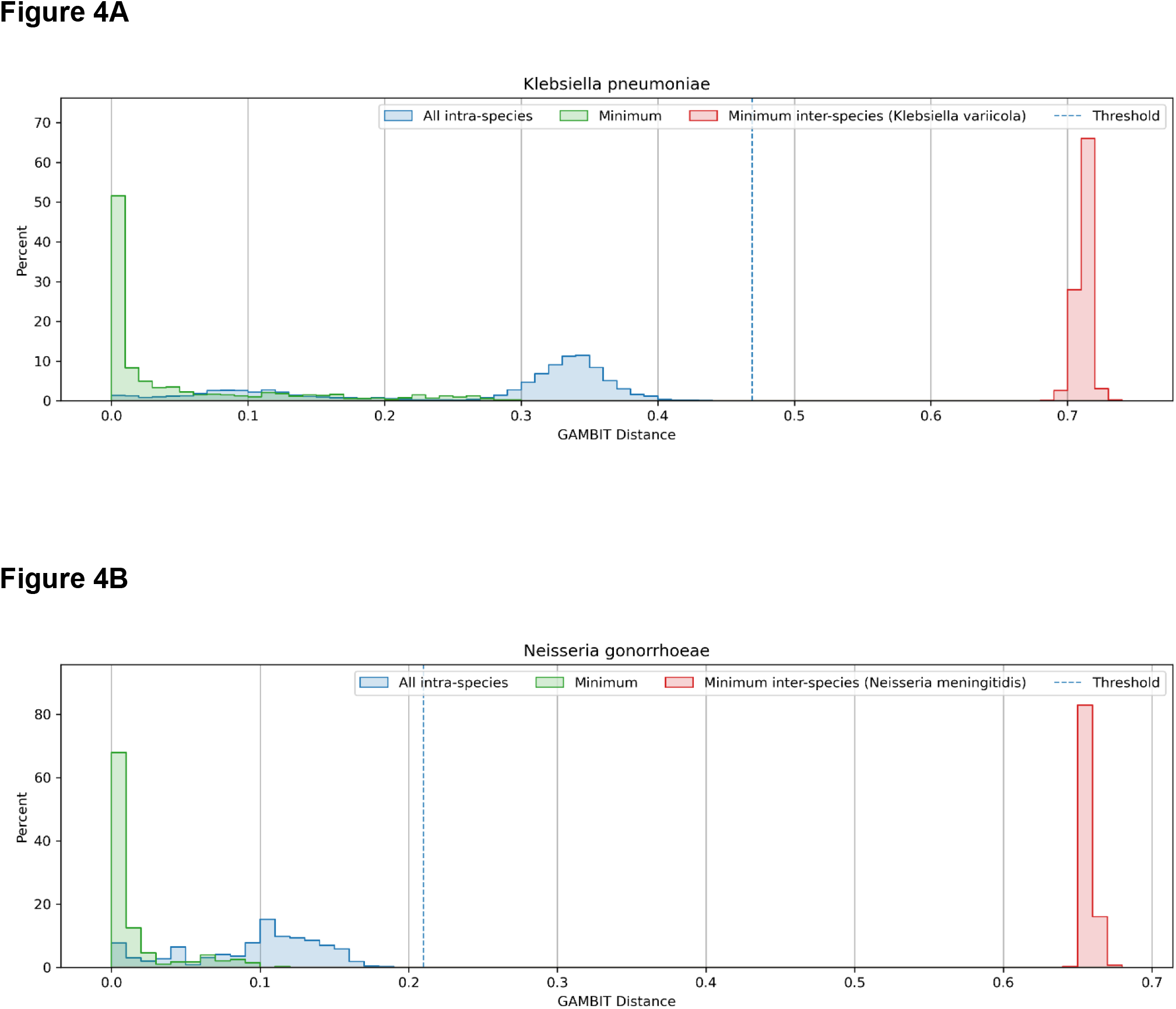
Distribution of GAMBIT distances within a species and to the nearest sister taxon. Three histograms are shown in each panel (normalized as fractions of total scores for comparison sake). The green histogram represents distribution of GAMBIT distances for the top hit for each isolate in the species. The blue histogram represents the distribution of GAMBIT distances for all pairwise comparisons within the species. The red distribution represents the distribution of GAMBIT distances for all pairwise comparisons between the species of interest and all isolates of that species closest sister taxon. The dashed blue line represents both the threshold for that species and the max intra GAMBIT distance. For panel A *Klebsiella pneumoniae* is shown (its closest sister taxon is *Klebsiella* variicola). For panel B *Neisseria gonorrhoeae* is shown (its closest sister taxon is *Neisseria meningitidis*).

Once these values are calculated we compare max intra to min inter for each species. For cases where max intra is less than min inter, we used max intra as the threshold (Figure 4 AB). Thus, no isolate in our reference database could generate a false positive with this threshold. The Jaccard distances for *Klebsiella pneumoniae* and *Neisseria gonorrhoeae* are plotted in Figure 4 as examples. There are 877 *Klebsiella pneumoniae* isolates in the GAMBIT database. In Figure 4A we plot three distributions of GAMBIT scores. The green distribution represents the top hit (min GAMBIT distance) for each of the 877 *Klebsiella pneumoniae* isolates in the database. This is relevant as GAMBIT assigns genus and species information based on the top hit in the database. The blue distribution is all pairwise comparisons of the 877 *Klebsiella pneumoniae* isolates (877^2^). The red distribution is all the pairwise scores between the 8777 *Klebsiella pneumoniae* isolates and its closest taxon by GAMBIT distance *Klebsiella variicola*. This figure illustrates how most species in our database are well separated by GAMBIT distance from their closest sister taxon. The max intra distance for *Klebsiella pneumoniae* is 0.469 (denoted by the dashed blue line in Figure 4A) and the min inter distance 0.689. These spreads give us confidence that our thresholds will not generate false positives. Figure 4B shows the same distributions for *Neisseria gonorrhoeae*. There are 208 isolates of *Neisseria gonorrhoeae* in the GAMBIT database. The max intra (and threshold) for *Neisseria gonorrhoeae* is 0.210 and the min inter to its closest sister taxon (*Neisseria meningitidis*) is 0.649. Of the 1,414 species in our database, we used this method to set the threshold for 1,371 of them (∼97%).

Next, we examined 25 species that have overlapping distributions of inter distances and intra distances. These species had only a minimal overlap in the distribution of the scores with none of the top hit distances near the min inter value. In these 25 cases, we used 95% of the min inter distance as the threshold. All of the top hits (which is used to assign species identification by GAMBIT) are well above the threshold. The goal of these thresholds is to empirically set a conservative threshold that if eclipsed allows GAMBIT to call with certainty identification at the species or genus level.

Lastly, there were 17 species that had more substantial overlap in their intra and inter distances with one or more sister species (Table 2). For these species we employed the strategy of dividing these species into between 2 and 5 subspecies groups based on clustering of GAMBIT distances. For these species GAMBIT is mapping to the subspecies level but reporting at the species level. At the subspecies level we were able to create thresholds based on either max intra or 95% min inter (Supplemental Table 4). An example of *Shigella boydii* is shown in Figure 5. *Shigella boydii* had overlaps with *Shigella dysenteriae* shown in the panel labeled combined (Figure 5A). However, after dividing *S. boydii* into two subgroups (Figure 5A), there is no overlap and clean thresholds can be established. The distribution of intra distances is shown in blue, while the distribution of inter distances with the overlapping species (in this case *S. dysenteriae*) is outlined in red. For the subgroups, the distribution of distances between that subgroup and the other subgroups of the species are shown in green. The threshold for each subgroup is shown with the vertical dashed blue line. Figure 5B demonstrates how we resolved the overlaps of *Pseudomonas amygdali* with *Pseudomonas savastanoi* (shown in purple) and *Pseudomonas syringae* (shown in red). Here we divided *Pseudomonas amygdali* into three subgroups, of which subgroups 1 and 3 had no remaining overlaps and subgroup 2 used the 95% of min inter distance to set the threshold.

**Table 2:** Species in the GAMBIT database that were broken into subspecies during the process of establishing thresholds.

| name | max intra | min inter | closest taxon | number of subgroups |
| --- | --- | --- | --- | --- |
| *Bacillus amyloliquefaciens* | 0.456449 | 0.34734836 | *Bacillus velezensis* | 2 |
| *Bacillus cereus* | 0.902063 | 0.5682511 | *Bacillus thuringiensis* | 4 |
| *Bacillus pumilus* | 0.878503 | 0.7945259 | *Bacillus safensis* | 2 |
| *Brucella suis* | 0.067703 | 0.022016378 | *Brucella canis* | 2 |
| *Clostridium botulinum* | 0.994353 | 0.8220584 | *Clostridium sporogenes* | 3 |
| *Enterobacter cloacae* | 0.66532 | 0.28938964 | *Enterobacter hormaechei* | 4 |
| *Enterobacter hormaechei* | 0.607138 | 0.28938964 | *Enterobacter cloacae* | 2 |
| *Escherichia coli* | 0.682538 | 0.36488694 | *Shigella sonnei* | 3 |
| *Prochlorococcus marinus* | 0.998967 | 0.99406457 | *Arcobacter butzleri* | 4 |
| *Pseudomonas amygdali* | 0.522444 | 0.2951006 | *Pseudomonas syringae* | 3 |
| *Pseudomonas putida* | 0.886751 | 0.8334536 | *Pseudomonas monteilii* | 3 |
| *Pseudomonas savastanoi* | 0.489886 | 0.29782194 | *Pseudomonas amygdali* | 2 |
| *Pseudomonas syringae* | 0.914961 | 0.2951006 | *Pseudomonas amygdali* | 5 |
| *Salinispora pacifica* | 0.804802 | 0.52663755 | *Salinispora oceanensis* | 3 |
| *Shigella boydii* | 0.296933 | 0.20483686 | *Shigella dysenteriae* | 2 |
| *Shigella dysenteriae* | 0.56022 | 0.20483686 | *Shigella boydii* | 2 |
| *Streptococcus pseudopneumoniae* | 0.846564 | 0.58063465 | *Streptococcus mitis* | 2 |

**Figure 5.**
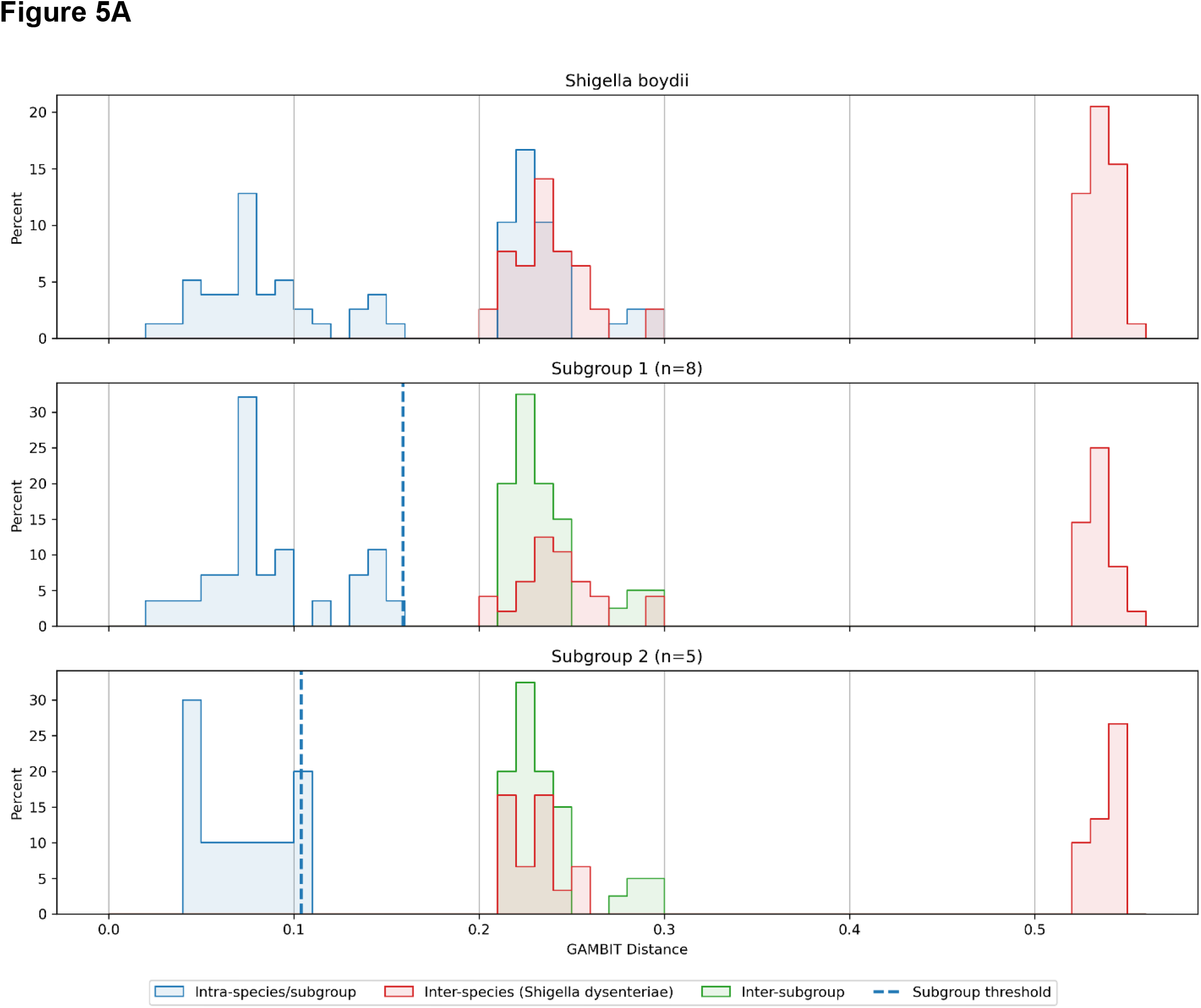
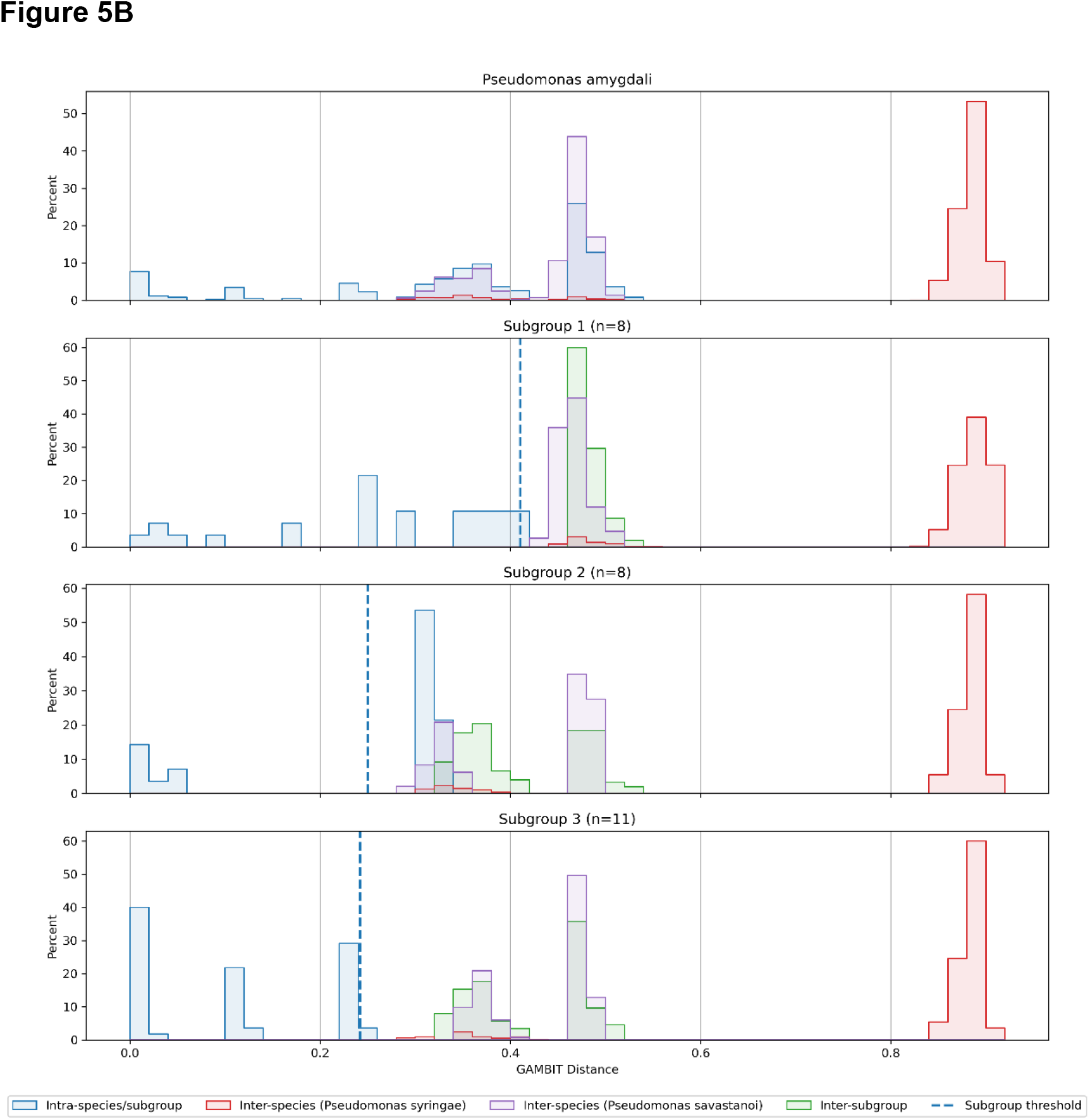
Threshold determination through subgrouping of species. A. *Shigella boydii*. The top panel shows the distribution of intra distances in blue and the distribution of inter distances with *Shigella dysenteriae* in red. The middle and bottom panels show the *Shigella boydii* subgroups 1 and 2 respectively. The distribution of subgroup distances are shown in blue, the distribution of inter distances with *Shigella dysenteriae* in red. The distribution of distance between subgroup 1 and subgroup 2 are shown in green with the subgroup threshold represented with a dashed blue line. B. *Pseudomonas amygdali*. The top panel shows the distribution of intra distances in blue, the distribution of inter distances with *Pseudomonas savastanoi* in purple and the distributions of inter distances with *Pseudomonas syringae* in red. The bottom three panels show the *Pseudomonas amygdali* subgroups 1, 2 and 3. The distribution of subgroup distances are shown in blue, the distribution of inter distances with *Pseudomonas savastanoi* in purple and the distributions of inter distances with *Pseudomonas syringae* in red. The distribution of distance between subgroup presented in the panel and the other 2 subgroups are shown in green with the subgroup threshold represented with a dashed blue line.

Two final notes on our database construction at the species level. We could not cleanly differentiate *Lacticaseibacillus casei* and *Lacticaseibacillus paracasei* at the species level. Thus, GAMBIT reports the result ‘*Lacticaseibacillus casei/paracasei*’ for top hits that map to the genomes for these two species. While GAMBIT can differentiate between the closely related organisms of Escherichia coli and the Shigella species, when the top hit for either of these does not meet the confidence threshold, GAMBIT reports back ‘*Escherichia coli/Shigella*’ rather than *Escherichia* or *Shigella* alone.

If the top hit distance for a new query in GAMBIT is larger than the species threshold, there is still a possibility that GAMBIT will make a call at the genus level. We repeated the same procedure to set thresholds for genus. 335 genera had no overlaps between their intra and inter distances, while 117 (∼26%) had overlaps and used 95% of min inter to set the thresholds. The total number of genera reported is two less than present in the database (452 vs 454) for two reasons. First, is the combining of ‘*Escherichia* and *Shigella* into a single reporting group as mentioned above. The second is *Eubacterium siraeum. Eubacterium* is a proposed genus not officially accepted in the NCBI taxonomy, so we only call *Eubacterium siraeum* at the species level.

### Validation of GAMBIT by Alameda County Public Health Labs

From 10/12/16 to 02/13/2020 the Alameda County Public Health Laboratory used whole genome sequencing utilizing GAMBIT for genus and species identification on 88 proficiency test samples. The source of these samples was either isolates identified at the Public Health Lab and confirmed by a reference laboratory; or they were specimens provided as quality assessment standards through proficiency testing (College of American Pathologists). Using the conservative thresholds described in the previous sections, GAMBIT reported zero false positives out of these 88 samples (Table 3 and Supplemental Table 5). GAMBIT predicted the correct genus or species needed to pass the proficiency test in 86 of the 88 samples. The other 2 samples were correctly predicted and reported using an *in silco* 16S test on the whole genome sequencing (see materials and methods for details). *In silco* 16S was used on an additional sample to report the species level information but only genus level information was expected for the proficiency test (GAMBIT reported *Pseudomonas* genus, 16S reported *Pseudomonas putida*). One of these samples (*Proteus vulgaris*) the top genus prediction for GAMBIT was correct but the scoring method did not pass the threshold. The only case where the top match was not a genus match was for *Granulicatella adiacens*. Neither this species nor any member of the genus *Granulicatella* are in the GAMBIT database and thus no correct match can be expected. In summary, in 87 out of 87 proficiency test samples where the species was present in the GAMBIT database the top match from GAMBIT provided the correct reporting information to satisfy the test. In the only instances the score associated with the top match did not exceed the threshold used by GAMBIT, the top match was correctly confirmed with 16S analysis. The *Granulicatella adiacens* had to be identified by 16S because the GAMBIT database did not contain any representatives from the genus *Granulicatella*.

**Table 3:** Summary of GAMBIT Validations from Alameda County Public Health Labs and Nevada State Public Health Labs.

| Validation Set | Isolates | Total Species | Total Genera | False Positives | Species-level calls | Genus-level Calls | No identification |
| --- | --- | --- | --- | --- | --- | --- | --- |
| Alameda County Public Health | 88 | 51 | 29 | 0 | 82 | 4 | 2 |
| Nevada State Public Health | 605 | 25 | 15 | 0 | 580 | 27 | 0 |

The 82 species predictions made by GAMBIT described above spanned 51 different species over 29 unique genuses. This included correct distinctions between 5 different species of *Staphylococcus* (*S. aureus, S. epidermidis, S. hominis, S. ludgunesis* and *S. saprophyticus*) and 8 different species of *Streptococcus* (*S. agalactiae, S. anginosus, S. constellatus, S. dysgalactiae, S. mutans, S. pneumoniae, S. pyogenes, S. salivarius*). Additionally, GAMBIT correctly differentiated one sample of *Shigella sonnei* from six other samples of *Escherichia coli*. The breadth of genus and species correctly predicted by GAMBIT combined with the discriminatory power within genuses such as *Staphylococcus* and *Streptococcus* demonstrates the utility of GAMBIT in clinical diagnostics.

### Validation of GAMBIT by Nevada State Public Health Labs

From 2021 to 2022 the Nevada State Public Health Lab (NSPHL) performed identifications on 605 isolates. In addition to performing bacterial identification with the normal procedures, all isolates were sequenced on an Illumina sequencing platform and identifications generated with GAMBIT. For this validation we compare GAMBIT identifications with the primary identification generated by the NSPHL. NSPHL is a reference laboratory and utilizes multiple methods of microbial identification, including biochemical methods, MALDI-TOF and genomic means (sequencing or probe-based methods). These additional methods were used to determine the identification of the isolate when the initial result was undetermined or called at the genus level only. Thus, NSPHL provides a “gold standard” identification with which to compare GAMBIT identifications.

For all 605 isolates, GAMBIT did not provide any incorrect identifications. This is by design, for the thresholds used by GAMBIT to be conservative and err on the side of genus only or no call rather than produce a false positive identification. Results of this validation are summarized in Table 3. For 518 isolates, GAMBIT and the initial NSPHL identification provided the same identification at the species level. These results spanned 25 species and 15 genera.

For 62 isolates, GAMBIT provided a more specific identification than the initial NSPHL identification (see Supplemental Table 6). In two instances, GAMBIT returned the correct genus where the initial NSPHL returned no ID. In another instance, GAMBIT returned the species where the initial NSPHL returned only the genus. There were 49 instances where GAMBIT correctly returned IDs for *Shigella flexneri* and *Shigella sonnei* where the initial NSPHL ID was *Escherichia coli*. The ability of GAMBIT to distinguish *Shigella* from *E. coli* with confidence is a compelling feature for public health applications. Lastly, there were nine instances where GAMBIT called a more specific species variant while the initial NSPHL ID was a more genetic species call, an example of which is GAMBIT calling *E. hormaechei* when the initial NSPHL ID was *E. cloacae*.

For 25 isolates GAMBIT provided only the correct genus where the initial NSPHL identified at the species level (Supplemental Table 6). These instances fell broadly into two classes. The first class is where GAMBIT routinely identifies the species well, but a particular isolate or subset of isolates were called at the genus level because of the conservative thresholds. For example, out of 209 *Klebsiella pneumoniae* in this validation set, GAMBIT called 208 at the species level and 1 at the genus level. 8 isolates fall into this category (*Klebsiella, Pseudomonas* and *Salmonella* genus only calls listed in Supplemental Table 6). The other class covering 17 isolates from the validation set represent species which are not well-represented in the GAMBIT database and thus are often called at the genus level. An example is that of 13 *Enterobacter cloacae* only 3 were called at the species level, the other 10 were called at the genus level. These include *Enterobacter cloacae, Mycobacterium chimaera* and *Camplylobacter lari* from this validation set. Future updates to the GAMBIT database could improve performance for these organisms.

### Tree-building tool based on GAMBIT distances

A benefit of our k-mer based method is the rapid generation of relative relatedness trees utilizing the Jaccard distances. A user can enter a number of genome assemblies from which they want to generate a relative relatedness tree. GAMBIT’s tree-building utility generates all pairwise comparisons amongst those genomes and stores them in a matrix. Hierarchical clustering is then performed on this matrix and a relative relatedness tree is quickly generated. The distance values generated by this process are highly correlated to sequence identity. Because only a subset of the genome is utilized to calculate the GAMBIT score, this relatedness method is inferior to methods that provide comparisons based on comparing entire genomes (such as Mauve alignments). However, the benefit of this method is the ability to rapidly generate a relatedness tree with dozens of input genomes—something impractical to do if using whole genome comparison methods.

To validate the usefulness of this tool, we generated a tree from 29 *E. coli* strains that had been characterized by the Stephens Lab^28^. All 29 strains had been assigned to one of five phylogroups based on molecular testing (either group A, B1, B2, D or F). In addition, MLST subtyping was done on each strain which is a more granular division than phylogroups and results in a strain number such as 1193. The results of the relatedness tree are shown in Figure 6. Also shown in Figure 6 is the heat map of GAMBIT distances for all the strains. It is on these scores that the clustering occurs to generate the tree. Phylogroups are designated by different colored labels and the letter associated with the phylogroup. In all cases members of the same phylogroup cluster together. Additionally, in all cases where strains share the same MLST, they form tight clusters together. There were four instances of pairs of strains with the same MLST and three instances of a trio of strains that have the same MLST (Figure 6). This extension of the GAMBIT scoring system provides a useful tool for researchers and diagnosticians interested in the relatedness of their bacterial isolates.

**Figure 6.**
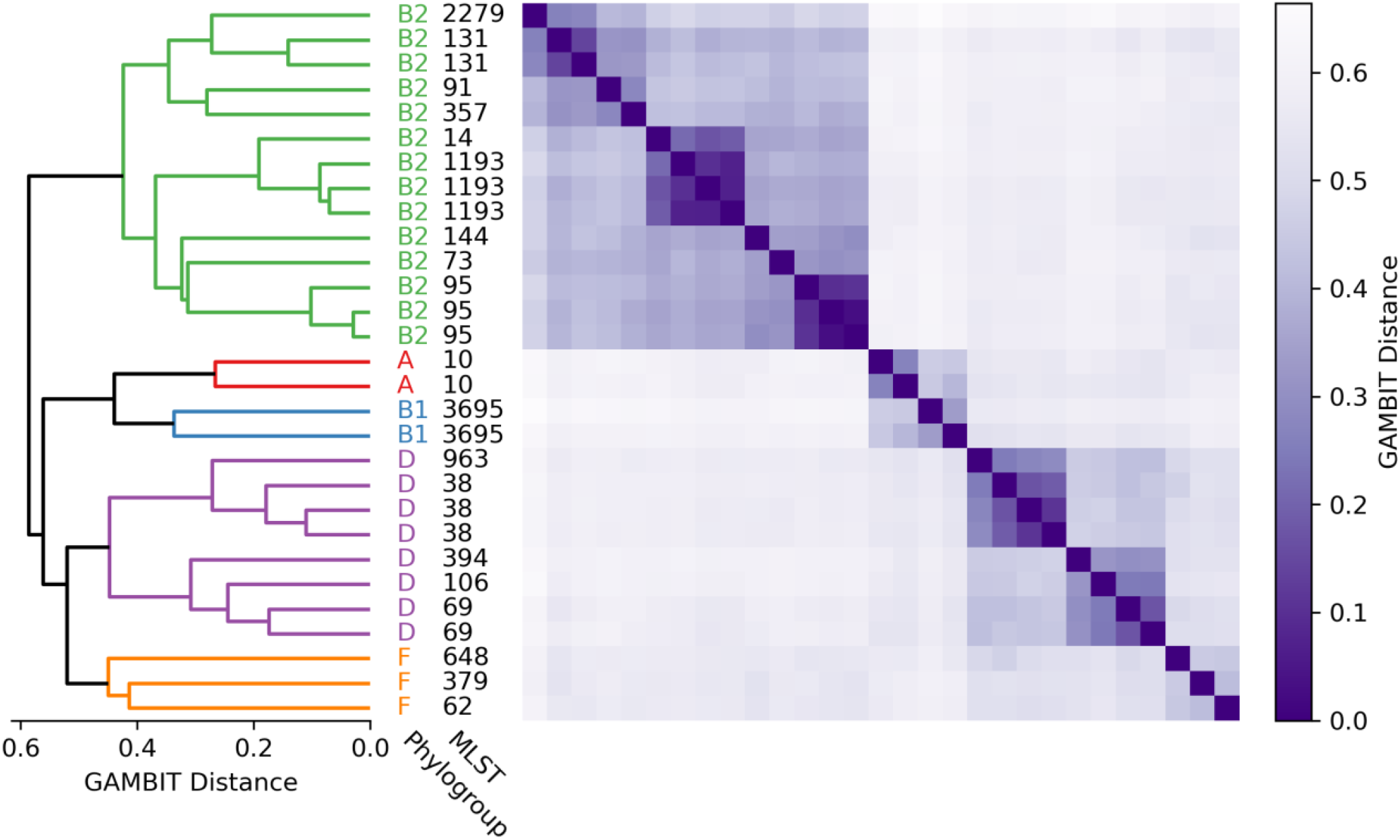
Phylogenetic tree estimated using GAMBIT distances. PCR and sequencing were used to perform multilocus sequence typing and determine the phylogroups of 29 *E. coli* genomes provided by the Stephens lab^27^. We then calculated all pairwise GAMBIT distances (displayed as a heatmap on the right of the figure) for these genomes using standard parameter values, and used the resulting distance matrix to perform hierarchical clustering using the UPGMA method. The resulting tree groups both phylogroups (colored) and MLST types into clades.

## Discussion

### Utilization of GAMBIT (and whole genome sequencing) as a lab developed test

Since its initial version of development, GAMBIT has been in use at the Alameda County Public Health Laboratory as a laboratory developed test (LDT). After extensive initial validation, the use of whole genome sequencing combined with GAMBIT analysis has been used in over 1500 instances of bacterial identification in a diagnostic setting. The vast majority of instances have resulted in outright identification while occasional instances required use of 16s rRNA or other sequence based analyses to accomplish complete identification. GAMBIT possesses certain features that make it functional in the diagnostic / CLIA-certified setting. Firstly, the database that is utilized for identification is highly curated. The curation process identified and removed several genomes in RefSeq that were either misidentified or mislabeled. Such genomes, had they remained in the database, might contribute to a lack of specificity for genome based LDT. Secondarly, the database is locally hosted, in the public health laboratory. This means that it meets the requirement of existing in a CLIA-certified environment where it can be part of an ongoing quality assurance program. Had the database existed off-site, it would have to exist in a CLIA-certified environment. This is because as the specificity component of the test, it plays a crucial role in execution of the test. Thirdly, GAMBIT provides actual and definitive “calls” of identification. Use of BLAST with online databases typically results in a line listing of suggested matches and associated probabilities. GAMBIT instead provides calls of identification based on quantitative overlap of shared kmers. This allows it to function with defined signal to noise parameters that lend extreme confidence to an identity call. Fourthly, GAMBIT provides an output which describes detailed final results and the quantitative aspects of those results that led to the identity call. This means that raw information regarding test performance is available for documentation of the test, which lends itself very well to quality control and quality assurance programs that are required of CLIA certification or other similar accreditation agencies.

The k-mer based algorithm that GAMBIT utilizes takes inspiration from the Aarestrup group^21^ and bears similarity to methods from the Phillippy group^23^. The focus for our group was to create a k-mer based tool that could be used diagnostically. To accomplish this we needed to solve two problems beyond the use of a k-mer based comparison to determine genome similarity. First, we needed a gold-standard reference database that was static so that the result of a query was not dependent on the point in time when the query was run. Public databases, such as Genbank, are constantly updated. Additionally, there is little to no curation of genomes and their species assignment in public databases. As such, they are not suitable for use in a diagnostic process. Additionally, k-mer comparisons rely on a significant amount of pre-processing of any reference library to achieve the performance gains in terms of speed. We accomplished this by curating and culling all bacterial genomes in RefSeq from date July 1^st^, 2016 resulting in the necessary static pre-processed database that represented 48,224 genomes from 1414 species. We believe this highly curated list of bacterial genomes with genus and species information will be of general interest to the scientific community. This database will be updated periodically to add new species and additional genomes to existing species.

The second challenge to diagnostic implementation of GAMBIT was developing a set of thresholds that allowed a confident genus and/or species prediction. This is in contrast to existing systems that rely on percent match data and ranking where no particular “call” is made. We leveraged our curated database to perform all pairwise comparisons within this database. Using these data, we developed thresholds based on the closest distance that any genome outside the species mapped to that species. We ensured that our method would not call a species if the distance of the inspected genome to the top hit in our database was not within that threshold distance. To date, in routine proficiency testing of GAMBIT, the algorithm has not resulted in a misidentification. We believe that this framework for determining thresholds will lend itself to future improvements by utilizing machine learning approaches to identify k-mers that are most associated with certain genus or species and giving more weight to those k-mers when calling a specific species.

### Benefits of Whole Genome Sequencing for Organismal Description and Molecular Epidemiology

While the use of whole genome sequencing for diagnostic microbiology may seem unnecessary, its routine use for this purpose is potent in many respects. 1. The use of sequencing eliminates the need for professionally trained, highly experienced bacteriologists to perform bacterial identification. This is important to consider in recognition of the amount of time/experience needed for skilled bacteriologists to be generated in a workplace. Fewer and fewer members of the diagnostic workforce are preparing for such careers. As such, this capability faces the need to be replaced by technology. 2. The use of genomic sequencing allows for the description of organisms in addition to their identification. This means that factors such as virulence and drug resistance can be ascertained immediately, along with genus and species. Intelligence gained in this manner provides sophistication to the medical response in a rapid manner. 3. That same descriptive power allows discrimination of organisms at high granularity, providing for subsequent molecular epidemiologic assessment. Multiple instances of certain species can immediately be ascertained for relatedness, allowing for the detection of outbreaks through diagnostic processes. Any organism identified in this manner can be a subject of “look back” to also allow elucidation of slowly developing outbreaks. 4. Genomic sequencing provides a view of organismal change over time. Normal bacterial diagnostics provide no insight into how organisms change over time which may be relevant to their threat level in public health or in the medical field. An example of this was seen with drug resistant *Neisseria gonorrhoeae*, whereupon whole genome sequences revealed that mating with additional *Neisseria* species, likely in the pharynx, was leading to genomic changes that had significant medical and public health implications^17^.

## Figure Legends

**Supplemental Figure 1.**
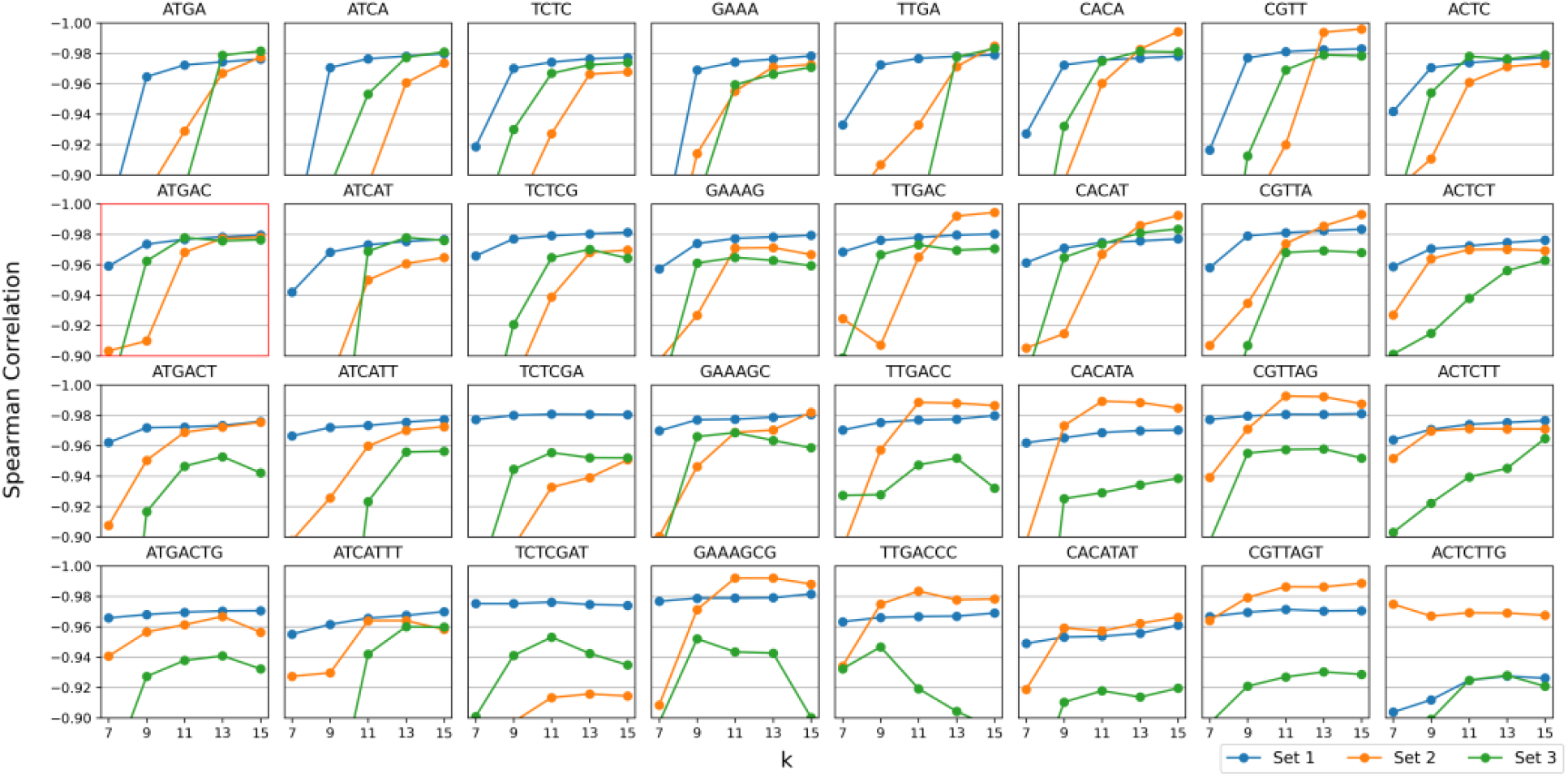
Spearman correlation of GAMBIT distance and ANI vs. value of *k* parameter for different choices of prefix. Each column corresponds to a different 7-mer base prefix. The first column uses our standard ATGAC prefix with two random nucleotides appended, the remaining seven were chosen randomly. Subplots within a column use this base prefix truncated to 4, 5, 6, or 7 nucleotides (the subplot using our standard prefix highlighted in red).

## Data availability

The 88 genomes from Alameda County Public Health Labs and the 605 genomes from the Nevada State Public Health Labs will be submitted to NCBI and linked to a single BioProject before final publication. The Snakemake workflow mentioned below can also be used to download the GAMBIT reference database and all genome assemblies used in this project.

## Code availability

GAMBIT source code has been made available through a publicly accessible GitHub repository: https://github.com/gambit/gambit/. The State Public Health Bioinformatics (StaPH-B) consortium maintains Docker container images of GAMBIT version releases accessible through the StaPH-B DockerHub and Quay repositories: https://hub.docker.com/r/staphb/gambit/ and https://quay.io/repository/staphb/gambit/, respectively. Dockerfiles for these images have been made available through the StaPH-B docker-builds GitHub repository: https://github.com/StaPH-B/docker-builds.

Additionally, WDL (workflow description language) workflows were developed by Theiagen Genomics to capture GAMBIT functionality and have been made available through the Theiagen Genomics Public Health Bacterial Genomics (PHBG) Dockstore Collection: https://dockstore.org/organizations/Theiagen/collections/PublicHealthBacterialGenomics/. While these WDL workflows can be run locally or on an HPC system at the command-line, utility was optimized for public health end-users with limited programming or bioinformatics experience through Terra, a bioinformatics web application developed by the Broad Institute of MIT and Harvard in collaboration with Microsoft and Verily Life Sciences: https://app.terra.bio/.

## Acknowledgements

This work was supported in part by the National Institutes of Health, grant R15AI130816-01A1. We thank the College of Arts and Sciences at Santa Clara University for supplemental funding.

